# Dynamic-ultrastructural cell volume (3D) correlative microscopy facilitated by intracellular fluorescent nanodiamonds as multi-modal probes

**DOI:** 10.1101/823278

**Authors:** Neeraj Prabhakar, Ilya Belevich, Markus Peurla, Xavier Heiligenstein, Huan-Cheng Chang, Cecilia Sahlgren, Eija Jokitalo, Jessica M. Rosenholm

## Abstract

Three-dimensional correlative light and electron microscopy (3D CLEM) are attaining popularity as a potential technique to explore the functional aspects of a cell together with high-resolution ultrastructural details across the cell volume. In order to perform such a 3D CLEM experiment, there is an imperative requirement for multi-modal probes that are both fluorescent and electron-dense. These multi-modal probes will serve as landmarks in matching up the large full cell volume datasets acquired by different imaging modalities. Fluorescent nanodiamonds (FNDs) are a unique nanosized, fluorescent, and electron-dense material from the nanocarbon family. We hereby propose a novel and straightforward method for executing 3D CLEM using FNDs as multi-modal landmarks. We demonstrate that FNDs is biocompatible and easily identified both in living cell fluorescence imaging and in serial block-face scanning electron microscopy (SB-EM). We illustrate the 3D CLEM method by registering multi-modal datasets.

## Introduction

Correlative light and electron microscopy (CLEM) combine the strengths of fluorescence and electron microscopy and it allows overcoming their respective limitations for cell imaging(1–3). CLEM can be employed to study dynamics and localization of macromolecules and proteins with live cell light microscopy (LM) followed by electron microscopic (EM) examination of the ultrastructural morphology of the specific cell of interest (4–9). Thus, functional and ultrastructural details of one cell are obtained by the integration of the two imaging modalities (10,11). To date, numerous experimental CLEM approaches have been reported(5,12–14). Apart from providing functional and ultrastructural information, recent CLEM methods have employed super-resolution fluorescence techniques to bridge the resolution gap between diffraction-limited fluorescence microscopy and EM(6,13,15–18). However, the majority of developed CLEM methods are based on the correlation of LM with 2D images of thin cell sections imaged with transmission electron microscopy (TEM)(16,17,19–22). Consequently, these CLEM methods provide very limited information on the z-axis direction, as TEM sections are generally restricted to slices of about 60-100 nm thickness and they may also be tilted relative to the image planes in the confocal image stack, resulting in uncertainty in the final correlation.

Considering the complex 3D organization of a cell, most of the critical 3D cellular information, especially in the z-direction, is generally under-explored. Therefore, 2D CLEM methods could be improved by employing instruments capable of performing 3D imaging (14,23–30) enabling CLEM methods correlating 3D information from both LM and EM. Combining 3D fluorescence microscopy with 3D EM would significantly improve the technical possibilities for investigating complex cellular processes across the full volume of a cell.

Recently, several volume-CLEM methods that demonstrate 3D correlation have been presented (31–36). Typically, there should be some common landmark to allow correlation, as 3D imaging from both LM and EM generates substantial image datasets. Preferably, these landmarks should be detectable with both modalities. Consequently, intracellular fluorescent and electron-dense landmarks are critical for the execution of 3D CLEM experiments. One such fluorescent and electron-dense CLEM marker is the fluorescent nanodiamond (FND)(19,37–39). (40–44). FNDs are non-toxic to cells, and being nanosized particles, they can be easily internalized in living cells via endocytosis(45–47). FNDs have excellent photostability, and they have non-blinking far-red emission, which makes them well-suited for imaging of living and fixed cells. We recently reported that FNDs are robust intracellular landmarks in 2D CLEM experiments(39).

In this article, a 3D CLEM method is demonstrated using 35 nm FNDs as intracellular landmarks for correlating cell volume datasets from live-cell confocal microscopy and serial block-face scanning electron microscopy (SB-EM).

## Material and Methods

### FND production

The synthesis and characterization of 35 nm FNDs have been previously reported(48). A brief synthesis protocol is presented as follows. Synthetic type Ib diamond powders with a nominal size of 100 nm (MDA, Element Six) were purified in acids and suspended in water. A thin diamond film of ~50 μm thickness were made by depositing the diamond suspension on a silicon wafer. The diamond film was then treated by a 3-MeV proton beam and nitrogen-vacancy defect centers were created by annealing the proton beam-treated nanodiamonds. To produce 35 nm FNDs, the 100 nm FNDs were first mixed with NaF powders and crushed together with a hydraulic oil press under a pressure of 10 tons. Smaller FNDs were isolated by centrifugation after dissolving the mixture in hot water to remove NaF.

### Cell culture

MDA-MB-231 cells (Human breast adenocarcinoma) were cultured in Dulbecco’s modified Eagle’s medium (DMEM) supplemented with 10% fetal bovine serum, 2mM L-glutamine, and 1% penicillin-streptomycin (v/v), over µ-Dish 35 mm ibidi gridded dishes (ibidi GmbH, Germany). 10 µg/ml of 35 nm FNDs particles were prepared in 1 ml of cell growth media. Then, the cell media with particles was added to the cells growing. The cells were allowed to incubate with FNDs for 24h. Staining with living cell dyes was performed as follows. The cells were washed three times with serum-free DMEM, after which 0.2 μl of Mitotracker (MitoTracker® Green, ThermoFisher Scientific Inc, USA) was first added to 1.5 ml of medium (without serum and antibiotics) and then drop by drop to the dish. MDA-MB-231 cells were incubated for 30 min at 37°C.

### 2D SEM

MDA-MB-231 cells (Human breast adenocarcinoma) were cultured in Dulbecco’s modified Eagle’s medium (DMEM) supplemented with 10% fetal bovine serum, 2mM L-glutamine, and 1% penicillin-streptomycin (v/v). 10 µg/ml of 35 nm FNDs particles were prepared in 1 ml of cell growth media. Then, the cell media with particles was added to the cells growing. The cells were allowed to incubate with FNDs for 24h. Cells were fixed with 5% glutaraldehyde s-collidine buffer, postfixed with 2% OsO4 containing 3% potassium ferrocyanide, dehydrated with ethanol, and flat embedded in a 45359 Fluka Epoxy Embedding Medium kit. Thin sections were cut using an ultramicrotome to a thickness of 100 nm. The sections were stained using uranyl acetate and lead citrate to enable detection with SEM. The Zeiss LEO 1530 (Zeiss, Germany) SEM instrument used was for imaging. The applied voltage was 15kV, the detector was the In Lens-detector. The secondary electron detector placed in the electron optics column.

### Confocal microscopy

The living cell 3D imaging was performed with a Leica TCS SP5 confocal microscope (Leica Microsystems, Germany), using a 63X oil objective. The cells were maintained at 37°C, 5% CO_2_ during the imaging. The MitoTracker® Green and the FNDs were excited by 488 nm argon laser. Fluorescence was collected at 510-550 nm and 650-730 nm with PMTs (Photomultiplier tubes) for MitoTracker® Green and FNDs respectively. The MitoTracker® Green was recorded in 3D stacks together with FND landmarks in living cells. After imaging, the cells were fixed with and sample preparation for SB-EM was performed.

### 3D SB-EM sample preparation

The specimens were prepared using a protocol modified from Deerinck et al. (2010)(49). The cells were fixed for 30 min at RT within a fixative mixture consisting of 2% glutaraldehyde, 2% PFA, 2 mM CaCl_2_ in 0.1 M NaCac buffer, pH 7.4. Washed five times with NaCac buffer containing 2 mM CaCl_2_. The cells were postfixed for 1 hour on ice bath in a fume hood with 2% OsO4 - 1.5 % K_4_[Fe(CN)_6_] - 2 mM CaCl_2_ in 0.1 M NaCac buffer, pH 7.4. The cells were washed 5 times with distilled water (DW). The cells were then incubated in 1% aqueous thiocarbohydrazide (TCH) for 10 min at RT. The cells were 5 times washed with DW. The cells were incubated in 1% OsO_4_ in DW for 30 min at RT. The cells were washed 5 times with DW. The cells were incubated with 1% uranyl acetate at +4°C overnight. Washed 5 times with DW at RT. Incubated in the pre-warmed lead aspartate solution at 60°C oven for 30 min. Washed 5 times with DW and followed by serial dehydration. The cells were dipped to an aluminum plate with resin-acetone solution containing acetone with 50% (v/v) Epon resin to incubate for 1h. Further, cells were incubated in 100% Epon resin, incubate 1 h RT. The cells were allowed to polymerize in an oven at 60°C for 28 h.

### 3D SB-EM Imaging

The area of interest with the selected cells was trimmed from the plastic block and mounted onto a pin using conductive epoxy glue (model 2400; CircuitWorks, Kennesaw, GA). The trimmed block was further trimmed as a pyramid and its sides were covered with silver paint (Agar Scientific Ltd, Stansted, UK). To improve conductivity the whole assembly was platinum-coated using Quorum Q150TS (Quorum Technologies, Laughton, UK). SB-EM data sets were acquired with a FEG-SEM Quanta 250 (Thermo Fisher Scientific, FEI, Hillsboro, OR), using a backscattered electron detector (Gatan Inc., Pleasanton, CA) with 2.5-kV beam voltage, a spot size of 2.9, and a pressure of 0.15 Torr. The block faces were cut with 50-nm increments and imaged with XY resolution of 25 nm per pixel. The collected 16-bit images were processed for segmentation using an open-source software Microscopy Image Browser(50) as follows: a) individual images were combined into 3D stacks; b) the combined 3D-stack was aligned; c) the contrast for the whole stack was adjusted and d) the images were converted to the 8-bit format.

### Image correlation

The multi-modal datasets were registered using the eC-CLEM plugin on the Icy bioimage analysis platform. In order to match the large datasets on a laptop (i7, 16Gb RAM memory), the EM stack was binned 4 times. The FM stack was matched to the binned dataset using the FNDs as landmarks, targeting the center of the FNDs aggregates both in LM and EM using orthogonal views from Icy. 9 FNDs were sufficient to achieve the good overlay accuracy depicted in this manuscript. Rigid registration was performed despite a recommendation by the software to apply for non-rigid registration. This decision was made after careful observation of the LM dataset. Since living cell imaging was performed on the 3D stack, the cell dynamics causes some of the FNDs to move during the image acquisition. This natural movement is uneven in all FNDs. Local inaccuracies in this registration are coherent with the cell movement observed. The weighing of each individual landmark operated by eC-CLEM compensate for the shifts observed between the LM and the EM dataset and rigid registration leads to an accurate full registration. To generate the final overlay, the transformation was applied to the LM dataset to match the original EM dataset using the “apply a reduced scaled transform to a full-size image” function from eC-CLEM (Advanced usage). This final overlay was used to generate the movies in supplementary data.

## Results

### FND facilitated 3D cell volume-CLEM

Our 3D CLEM workflow begins by seeding FND incubated MDA-MB-231 cells over gridded glass-bottom dishes designed for CLEM experiments. Two FND incubated MDA-MB-231 cells (Figure. 1a) were selected for the 3D CLEM experiment. The usefulness of FNDs in the 3D CLEM experiment was demonstrated by staining MDA-MB-231 cells with a mitochondrial marker dye MitoTracker (Figure. 1b-c).

**Figure 1.**
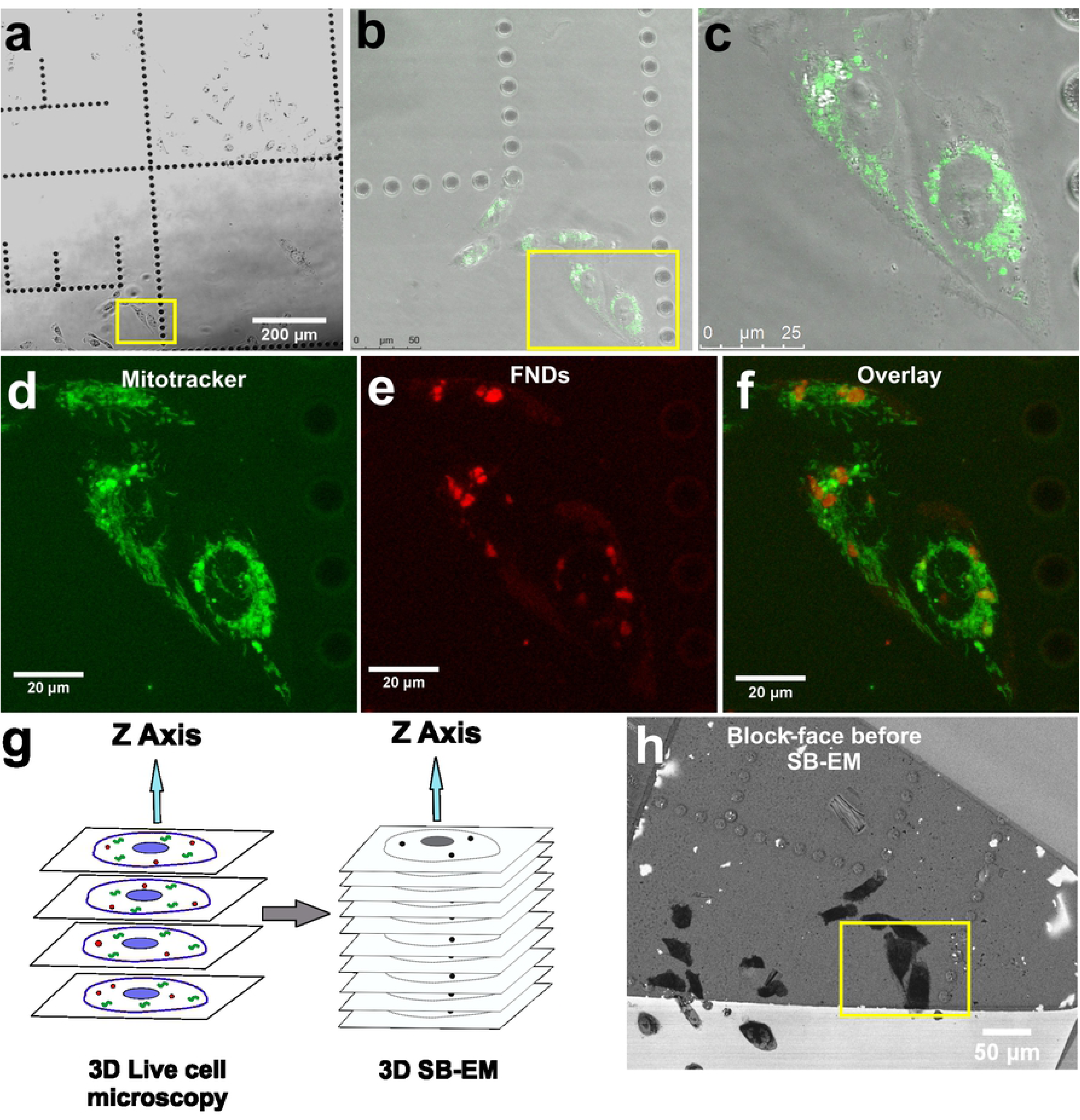
The workflow of the live cell confocal – SB-EM 3D CLEM experiment. a) Brightfield image of the living cells on a gridded glass-bottom dish. The cells selected for 3D CLEM are indicated by the yellow box. b) MitoTracker signal from the stained selected cells (indicated by the yellow box). c) Close up view of the selected cells shown by overlay of brightfield and MitoTracker images. d) Maximum intensity z-projection image from MitoTracker channel of the selected cells. e) Maximum intensity z-projection image from FND channel of the selected cells. f) Overlay of maximum intensity z-projections of MitoTracker and FND channels. g) Schematic representation of 3D CLEM workflow with FNDs (red dots) and a standard organic fluorophore (green structures). h) Localization of the selected cells on the EM block-face (near the letter E) for SB-EM. The yellow box indicates the same selected cells as in a).

Confocal image stacks of the whole cell volumes were acquired from both the MitoTracker (green) and the FND (red) signals (Figure. 1d-f and **Video. 1**). MitoTracker signal was seen widespread in the cytoplasmic space (Figure. 1d). The fluorescence signal from FNDs (Figure. 1e) was mostly localized to a few spots suggesting their confinement in vesicles in accordance with earlier results(39) FNDs are internalized by clathrin-mediated endocytosis (46,51) and they have a tendency to aggregate inside endosomal vesicles **(Figure. S1)** and subsequently slowly exocytose from cells(45,47). The aggregation of FNDs in cellular vesicles brings added benefit from a CLEM perspective(39) because, in comparison to single FNDs, the high concentration of FNDs aggregated inside vesicles provides better contrast both in fluorescence microscopy and in EM. In addition, confinement of FNDs in vesicles prevents their movement in the sample processing steps after confocal imaging enabling more reliable correlation of EM images.

3D localization of FNDs with respect to the MitoTracker fluorescence signal can be seen in **Video. 2**. These 3D confocal datasets were used for software-based correlation with SB-EM datasets. After confocal imaging, the selected cells were fixed, stained and embedded for SB-EM (Figure. 1g). The use of gridded glass-bottom dishes allowed easy identification of the cells of interest within the plastic block and trimming the blocks accordingly. The trimmed area was mounted on a pin and imaged in SEM. The mounted block-face overview image before SB-EM is displayed in Figure. 1h.

The two selected cells were identified (Figure. 1h) using a 15kV electron beam. The collection of 3D EM data was performed with an SEM instrument equipped with a system for serial block-face SEM (SB-EM). In SB-EM, an ultramicrotome performed automated sectioning of whole-cell volume by cutting thin sections (≥50 nm) from the sample’s block-face **(Video. 3)**. Consequently, after each cut, a high-resolution image of the freshly made block-face was acquired using a backscattered electron detector to form a 3D image stack. SB-EM imaging provided a three-dimensional dataset of the selected cells with a resolution to recognize the structure of interest (mitochondria) and FNDs aggregated in vesicular structures for CLEM (Figure. 2).

**Figure 2.**
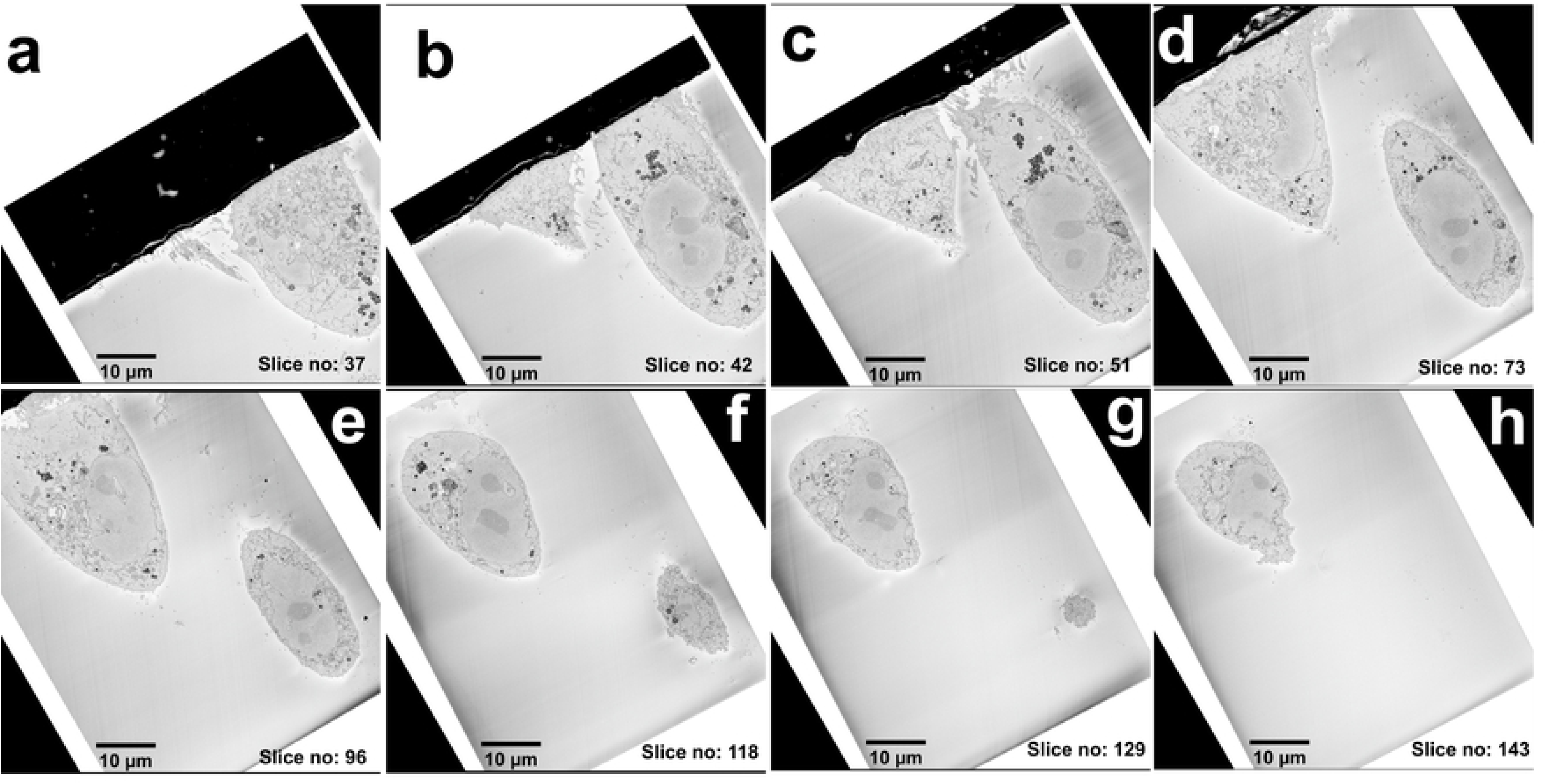
A collage of multi-plane images from the SB-EM 3D series. a-h) Images from the 3D EM stack acquired from the selected cells shown in the increasing order of z-axis. The distance between the slices is 50 nm.

Correlation of the LM and SB-EM volume datasets was done using the eC-CLEM plugin on the Icy bioimage analysis platform(52,53). First corresponding intracellular FNDs were identified in both datasets. FNDs aggregated in vesicles have a distinct appearance in SEM images (FigS1) and they are easily distinguished from morphological features of the cell. Figure 3 shows representative SB-EM and 3D LM image pairs (Figure. 3a and b; Figure. 3d and e) in which the corresponding FNDs are identified and marked.

**Figure 3.**
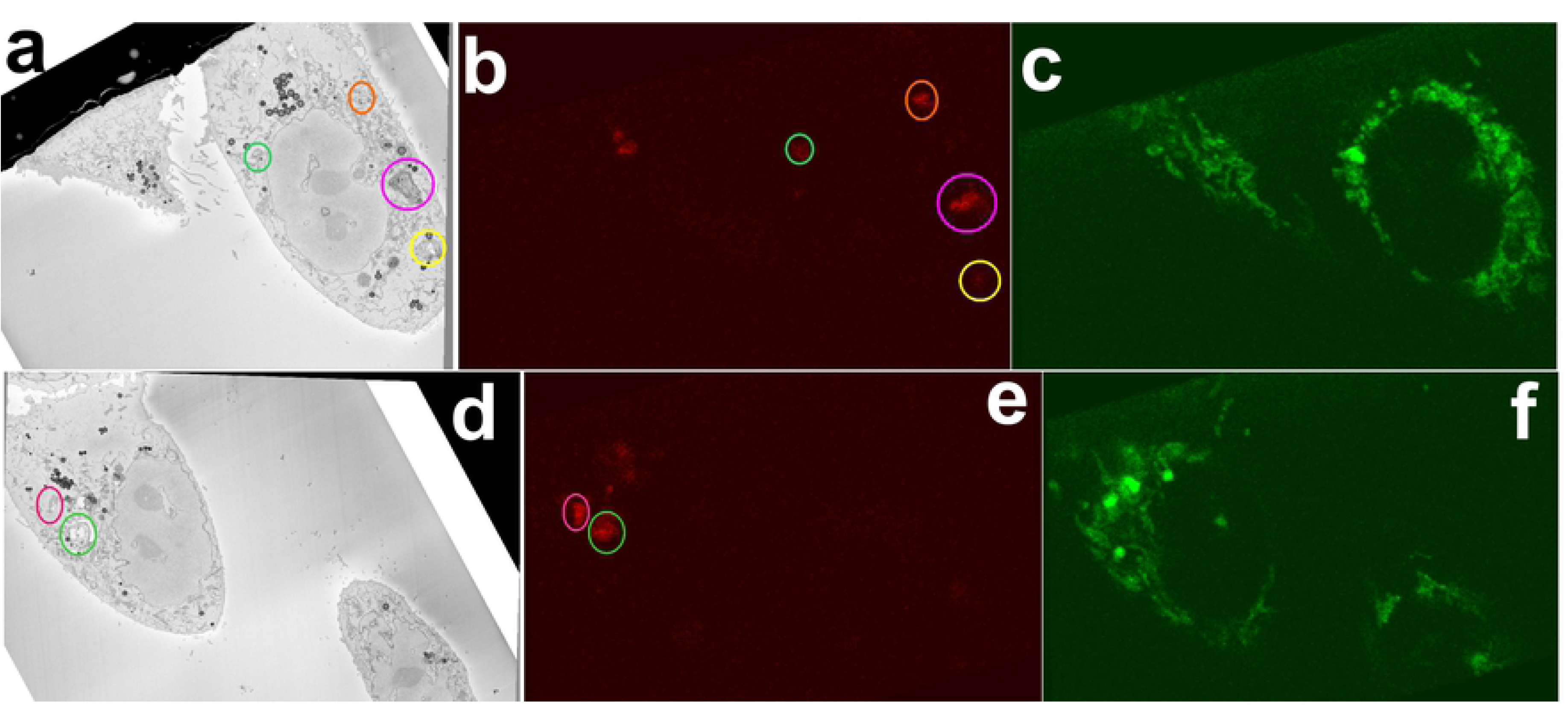
Overview of the corresponding FND aggregates in EM and LM. Corresponding SB-EM images (a and d) were matched in respective with FND (color-coded) localized in vesicles (b and e). Corresponding MitoTracker channels are shown in c and f.

Correlation of the two volume datasets was calculated using the identified FND position pairs as fiducials, and the accordingly transformed volume dataset of the fluorescence signal of interest (MitoTracker) was overlayed on the SB-EM stack (Figure. 4).

**Figure 4.**
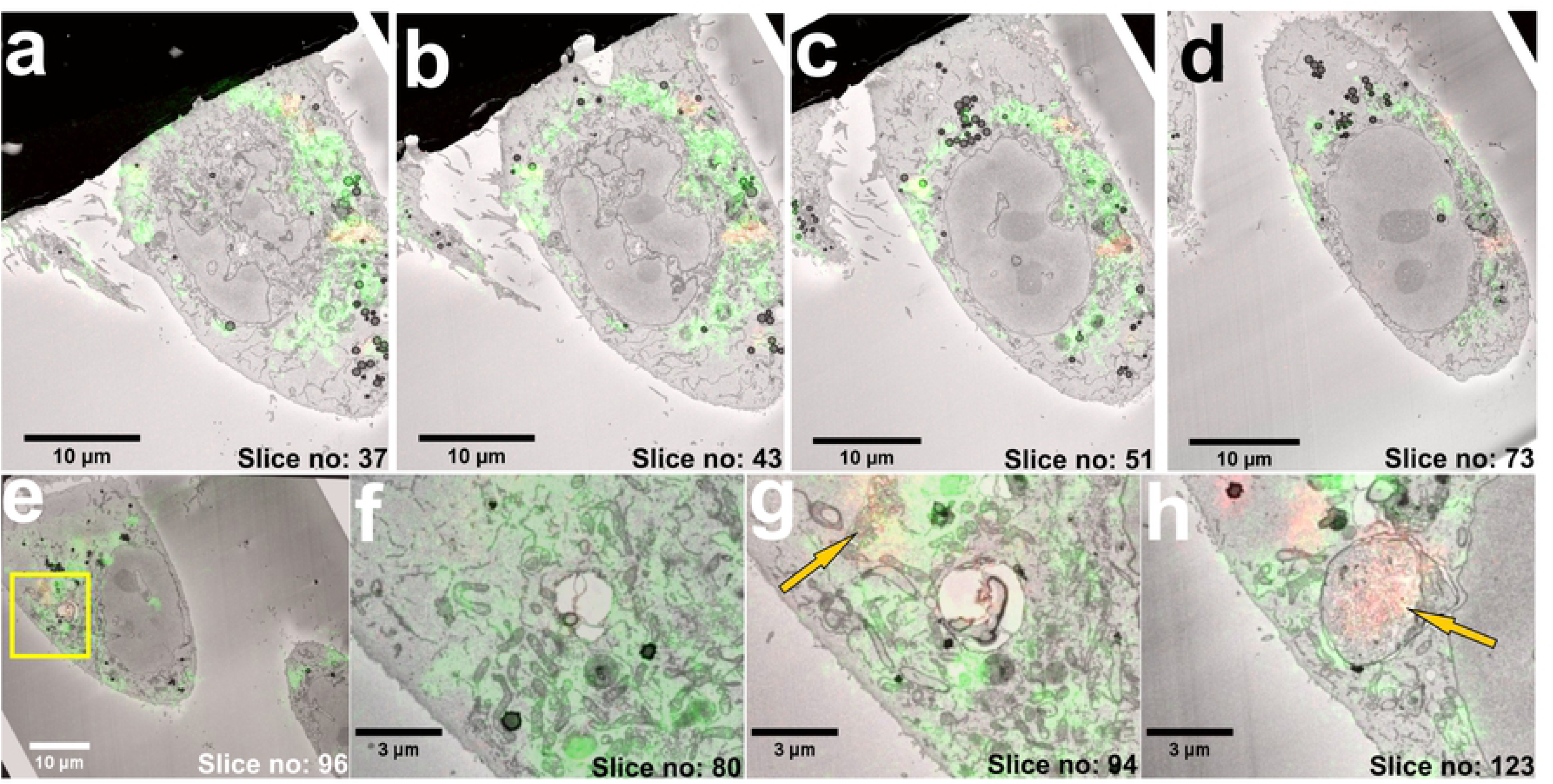
Volume-CLEM of living cells. a-d) CLEM images of multiple planes in increasing z-axis. e) ROI selected in CLEM image to demonstrate the correlation of LM over EM. f) In slice no:80, a single high-resolution image shows numerous mitochondria (green) and an empty vesicle. g) In slice no: 94, FNDs (red), mitochondria (green) and a vesicle can be seen. h) In slice no: 123, a vesicle filled with FNDs (red), and fewer mitochondria (green) can be seen. FNDs in the EM image are the dark dots inside the vesicle.

The FNDs facilitated mapping of the mitochondrial locations throughout the cell volume (Figure. 4a-d) resulting in good colocalization of MitoTracker signal with mitochondria seen in the EM image stack (**Video. 4-6 and Figure. S2**).

Quite commonly in literature the multi-modal correlation is performed in the absence of such a common landmark, and the process of 3D dataset correlation severely suffers from misalignment and errors in localizing critical information across 3D. However, in this type of CLEM approach, there can be multiple factors that could affect the precise image correlation. The major challenge encountered in the CLEM experiment was inherently low axial (600 nm) and lateral (250 nm) resolution provided by confocal microscopes compared to the nanometer scale resolution provided by EM. Currently, the limited resolution of confocal microscopy can originate in misalignment of details within large scale datasets. Sample autofluorescence, unspecific binding of fluorophores and obtaining a bright FND signal with live cell imaging are additional parameters still have to be optimized.

## Discussion

We have introduced a novel FND enabled cell volume (3D) correlative microscopy method. The CLEM workflow is straightforward and can be performed without any dedicated CLEM imaging systems. We demonstrated that a standard organic fluorophore can be used for 3D CLEM experiments with the FND based method without any special sample preparation requirements. In general, organic fluorophores do not survive routine EM sample processing and are not electron dense molecules, and therefore are not detectable with EM. In contrast, the employed 35 nm FNDs were intracellularly detectable with both imaging modalities in our experiments, enabling successful correlation of volume datasets for 3D CLEM. FNDs can offer multiple advantages over currently used CLEM fiducials as their internalization does not need chemical permeabilization, which has impacts on cellular morphology and ultrastructure. FNDs may be considered as a leading contender in the search of an exceptional CLEM probe because they are not prone to chemical degradation, have excellent photostability, and their nanoscale size facilitates their rapid internalization to cells. In our CLEM workflow, confocal microscopy was chosen for the 3D living cell imaging even if it offers limited resolution. Pairing confocal with SB-EM imaging was a practical choice for our experiment because the specific instrument was available to us. However, the focused ion beam imaging (FIB-SEM) could be used as an alternative for automatically obtaining the serial section image stacks. However, SB-EM can manage larger sample volumes than FIB-SEM, but with more limited z resolution. Our next step is to explore the possibilities of performing FND enabled CLEM with 3D super-resolution imaging.

## Author contribution

The manuscript was written through contributions of all authors. All authors have given approval to the final version of the manuscript. N.P. designed the study and performed live cell microscopy.

I.B. performed the 3D SB-EM. M.P. performed SB-EM cell sample preparation. X.H. performed image correlation. H-C.C. supervised the synthesized of FNDs. C.S., E.J and J.R. supervised the study and provided guidelines.

## Acknowledgments

The authors would like to acknowledge Jenni Laine and Kai-Lan Lin (Electron microscopy unit, University of Turku) for providing technical assistance with sample preparation for SEM and Mervi Lindman, University of Helsinki for excellent technical assistance in the preparation of SB-EM specimens. Helen Cooper is acknowledged for kindly providing mitochondrial staining reagents. Financial contribution from Academy of Finland (Project Nos. 309374) is also greatly acknowledged. The authors acknowledge Biocenter Finland for financial support and Euro-BioImaging (www.eurobioimaging.eu) for providing access to imaging technologies and services via the Finnish Advanced Light Microscopy Node (Helsinki, Finland). We also thank Fen-Jen Hsieh for the preparation of 35 nm FNDs.

